# A role for Gic1 and Gic2 in Cdc42 polarization

**DOI:** 10.1101/407569

**Authors:** Christine N. Daniels, Trevin R. Zyla, Daniel J. Lew

**Affiliations:** Department of Pharmacology and Cancer Biology, Duke University, Durham, NC, USA

## Abstract

The conserved Rho-family GTPase Cdc42 is a master regulator of polarity establishment in many cell types. Cdc42 becomes activated and concentrated in a region of the cell cortex, and recruits a variety of effector proteins to that site. In turn, many effectors participate in regulation of cytoskeletal elements in order to remodel the cytoskeleton in a polarized manner. The budding yeast *Saccharomyces cerevisiae* has served as a tractable model system for studies of cell polarity. In yeast cells, Cdc42 polarization involves a positive feedback loop in which effectors called p21-activated kinases (PAKs) act to recruit a Cdc42-directed guanine nucleotide exchange factor (GEF), generating more GTP-Cdc42 in areas that already have GTP-Cdc42. The GTPase-interacting components (GICs) Gic1 and Gic2 are also Cdc42 effectors, and have been implicated in regulation of the actin and septin cytoskeleton. However, we report that cells lacking GICs are primarily defective in polarizing Cdc42 itself, suggesting that they act upstream as well as downstream of Cdc42 in yeast. Our findings suggest that feedback pathways involving GTPase effectors may be more prevalent than had been appreciated.

## Introduction

Regulation of cell shape is central to cell proliferation as well as many aspects of cell function. Cell shape is in large part governed by the cytoskeleton, which itself is regulated by multiple signaling pathways. Among the most prominent and ubiquitous cytoskeleton-regulating pathways are those mediated by evolutionarily conserved small GTPases of the Rho family, including Rho, Rac, and Cdc42 (Hall et al., 1993). These GTPases are thought to act as molecular switches, toggling between an inactive GDP-bound state and an active GTP-bound state. Intrinsic rates of activation (GDP/GTP exchange) and inactivation (GTP hydrolysis) are slow, and can be greatly enhanced by guanine nucleotide exchange factors (GEFs) and GTPase activating proteins (GAPs), respectively (Hodge and Ridley, 2016). Rho-family GTPases are prenylated and reside primarily on the cytoplasmic leaflet of cellular membranes, although they can be extracted to the cytoplasm by guanine nucleotide dissociation inhibitors (GDIs) (Boulter et al., 2010; Mitin et al., 2012). Signaling pathways controlling cell shape often act by regulating and localizing the activities of GEFs and GAPs, leading to specific spatiotemporal patterns of GTPase activity.

Information encoded by the abundance and spatial pattern of GTPase activity is decoded by a set of GTPase-specific “effectors”, which are proteins that bind to the active but not the inactive form of the GTPase. Most known effectors are cytoplasmic proteins whose activity and localization within the cell can change as a result of GTPase binding. Effector localization and activity can also be regulated by other signals (e.g. phosphoinositides), allowing for complex combinatorial control of the cytoskeleton. Among the most intensively studied effectors are the p21-activated kinases (PAKs)(Rane and Minden, 2014), the WASP and WAVE regulators of branched actin nucleation by Arp2/3 complexes (Alekhina et al., 2017), and the formins that nucleate and accelerate polymerization of unbranched actin filaments (Kovar, 2006). In aggregate, GTPase signaling via effectors is responsible for sculpting the cytoskeleton.

One major role for Cdc42 and Rac concerns the establishment of cell polarity (Etienne-Manneville, 2004). Studies of polarity establishment in the model yeast *Saccharomyces cerevisiae* led to the identification of both positive feedback and negative feedback loops built into the polarity circuit (Chiou et al., 2017; Howell et al., 2012). In the positive feedback loop, effector PAKs are recruited to bind GTP-Cdc42, and they bind a scaffold protein called Bem1, which in turn binds to Cdc24, the yeast GEF for Cdc42 (Kozubowski et al., 2008). These interactions mean that wherever there is a slight local accumulation of GTP-Cdc42, recruitment of PAK-Bem1-Cdc24 will lead to enhanced GEF activity, leading to further local Cdc42 activation in a positive feedback loop (Johnson et al., 2011). Once GTP-Cdc42, PAKs, and Cdc24 co-accumulate to high levels due to positive feedback, the active PAKs promote multi-site phosphorylation of Cdc24 (Bose et al., 2001; Gulli et al., 2000; Wai et al., 2009). This phosphorylation reduces GEF activity (Kuo et al., 2014), possibly by more than one mechanism (Rapali et al., 2017), yielding a negative feedback loop. Thus, in addition to signaling to the cytoskeleton downstream of the GTPase, some effectors can also act as feedback transducers to regulate the local activation of the GTPase itself.

Analysis of several Cdc42 and Rac effectors, including the PAKs, led to the identification of a conserved Cdc42/Rac interactive binding (CRIB) motif that recognizes GTP-Cdc42 and GTP-Rac (Burbelo et al., 1995). Bioinformatic searches for other CRIB-containing proteins identified the GTPase interacting components (GICs), Gic1 and Gic2, in *S. cerevisiae* (Brown et al., 1997; Chen et al., 1997). GICs are small proteins that encode membrane-binding amphipathic helices (Takahashi and Pryciak, 2007) and a short conserved GIC motif of unknown function (Jaquenoud and Peter, 2000) in addition to the CRIB domain. The mammalian binder of Rho GTPase (BORG) proteins have a similar organization and may constitute homologs of the GICs (Joberty et al., 1999). In yeast cells, GICs are concentrated at polarity sites marked by active Cdc42 (Brown et al., 1997; Chen et al., 1997). Deletion of either *GIC1* or *GIC2* does not produce a dramatic phenotype, but cells lacking both GICs are large and misshapen, and (in haploids) fail to proliferate at high temperature (37°C)(Brown et al., 1997; Chen et al., 1997).

Subsequent work implicated GICs in regulating both the actin and septin cytoskeletons. In yeast cells, filamentous actin is present in actin cables (linear filament bundles oriented towards the polarity site that enable type V myosin-mediated cargo delivery to the bud) and in cortical actin patches (branched actin structures that promote invagination of the plasma membrane at sites of endocytosis)(Kaksonen et al., 2006; Pruyne et al., 2004). In polarized cells, actin patches accumulate near the polarity site and cables are oriented towards that site. However, in *gic1*Δ *gic2*Δ haploids at 37°C, most cells display randomly distributed actin patches, and fail to form a bud (Chen et al., 1997). Moreover, Gic2 interacts with and helps to localize the formin Bni1 to the polarity site, providing a potential mechanism for actin regulation (Chen et al., 2012).

Septins are conserved filament-forming proteins that assemble into a ring surrounding the polarity site following polarity establishment in yeast (Gladfelter et al., 2001; Oh and Bi, 2011). However, in *gic1*Δ *gic2*Δ haploids at 37°C, most cells fail to recruit septins to the polarity site (Iwase et al., 2006). GICs were shown to bind septins and affect interactions between septin polymers in vitro, providing a potential mechanism for septin regulation (Iwase et al., 2006; Sadian et al., 2013).

In addition to the studies implicating GICs as mediators of Cdc42-induced actin and septin rearrangements, a genetic interaction was identified between GICs and the Ras-family GTPase Rsr1 (Kawasaki et al., 2003). Rsr1 mediates communication between various transmembrane “landmark” proteins, which mark preferred sites for subsequent polarization, and the Cdc42-based polarity establishment pathway (Howell and Lew, 2012). Unlike *gic1*Δ *gic2*Δ or *rsr1*Δ mutants, which are viable at 24°C, *gic1*Δ *gic2*Δ *rsr1*Δ triple mutants were lethal. This suggested that GICs might act upstream of Cdc42, in parallel with Rsr1, as well as downstream of Cdc42.

Here, we have investigated the *gic1*Δ *gic2*Δ phenotype in greater detail, using live-cell imaging of cells bearing probes for polarity regulators Cdc42 and Bem1. We found that at 37°C, a majority of *gic1*Δ *gic2*Δ cells failed to polarize at all. This finding provides an alternative interpretation for previous findings in which the mutants failed to polarize actin or septins: these defects could be secondary effects stemming from a more fundamental lack of Cdc42 polarization. A subset of the *gic1*Δ *gic2*Δ did polarize Cdc42 and Bem1 at 37°C, and those cells did not display any obvious difficulty in forming a bud, suggesting that downstream cytoskeletal defects (if present) were quite mild. We conclude that, as suggested by Kawasaki et al. (Kawasaki et al., 2003), a major role of the GICs is to promote Cdc42 polarization.

## Results

Polarity establishment in yeast is regulated by the cell cycle. In particular, activation of G1 cyclin-dependent kinase (CDK) complexes at a commitment point called start in G1 promotes Cdc42 polarization (Gulli et al., 2000). G1 CDK activation at start occurs through a transcriptional positive feedback loop in which rising CDK activity promotes the inactivation and nuclear export of the repressor Whi5, allowing more transcription of G1 cyclins (Costanzo et al., 2004; de Bruin et al., 2004; Skotheim et al., 2008). Commitment to enter the cell cycle (i.e. start) occurs when 50% of nuclear Whi5 has been exported, at which point the positive feedback loop becomes self-sustaining (Doncic et al., 2011). We used Whi5-GFP or Whi5-tdTomato as a probe for start, and Bem1-tdTomato (Howell et al., 2012) or GFP-Cdc42 (Kuo et al., 2014) as a probe for polarization. Haploid cells were grown at 24°C and arrested in G1 by treatment with mating pheromone. Arrested cells were released to proceed into the cell cycle by washing out the pheromone, and placed on microscope slabs at 37°C. Live cell imaging by confocal fluorescence microscopy was then employed to monitor probe localization.

In wild-type cells under these conditions, Whi5 nuclear export was closely followed by Bem1 (Fig. 1A, video 1) or Cdc42 (Fig. 1B, video 2) polarization. However, in *gic1*Δ *gic2*Δ cells there was a heterogeneous phenotype: a majority of cells failed to polarize either Bem1 (Fig. 1A, video 1) or Cdc42 (Fig. 1B, video 2) after Whi5 nuclear exit. A substantial minority of cells did polarize the probes, although polarity establishment occurred somewhat later than in wild-type cells. Quantification revealed that 30%-40% of *gic1*Δ *gic2*Δ cells were able to form buds, but compared to wild-type cells the start-to-budding interval was longer (Fig. 2A). Similarly, 30%-40% of *gic1*Δ *gic2*Δ cells were able to polarize Bem1 or Cdc42, but with a longer and more variable interval between start and polarization (Fig. 2B). For the subset of *gic1*Δ *gic2*Δ cells that did polarize, the interval between polarization and budding was similar to that in wild-type cells (Fig. 2C). Thus, the major defect exhibited by *gic1*Δ *gic2*Δ cells at 37°C was an inability to establish polarity.

**Figure 1:**
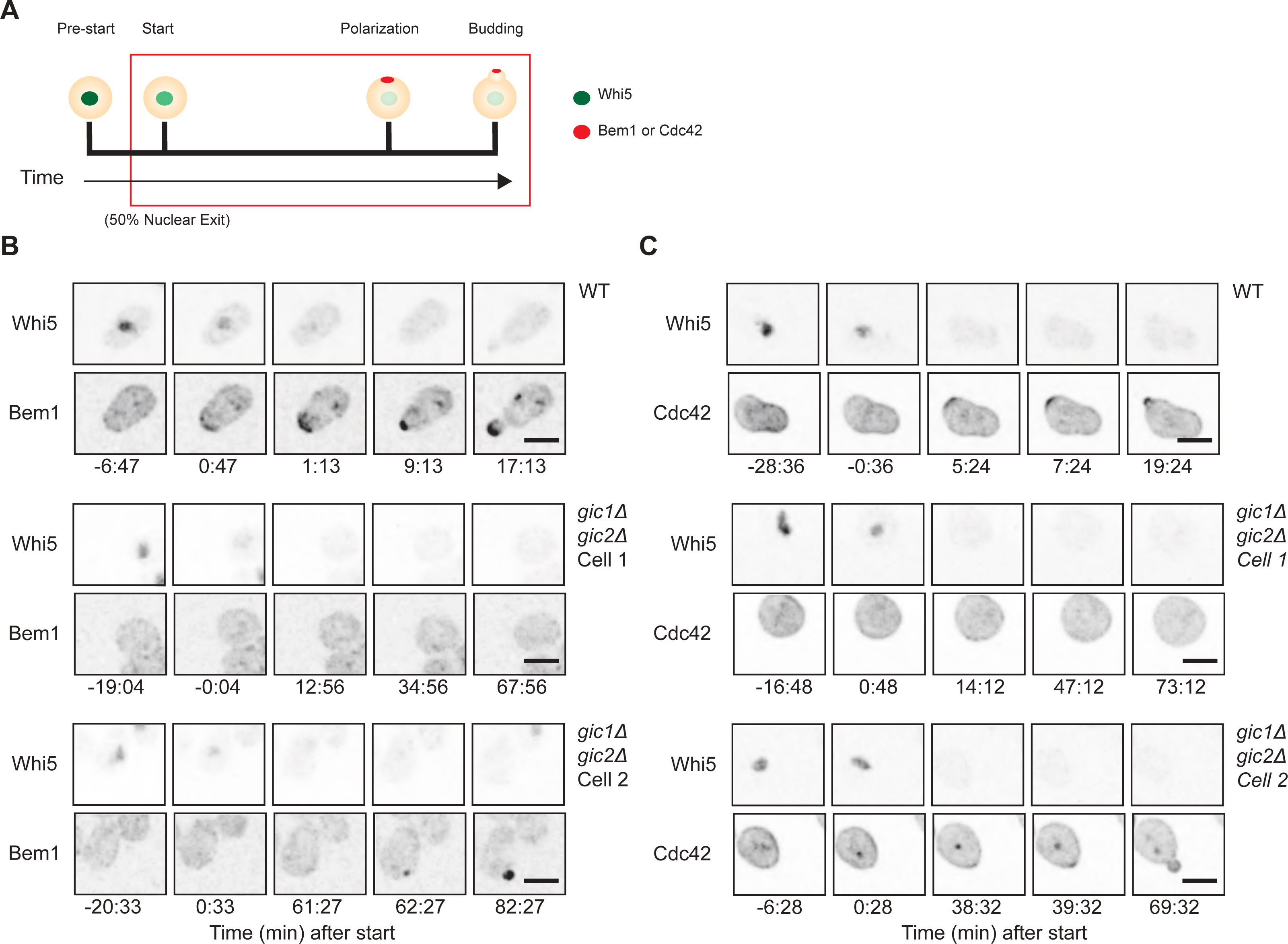
Delayed or blocked polarity establishment in gic1Δ gic2Δ mutants at 37°C. **(A)** Schematic depicting Whi5 and polarity protein distributions as cells proceed through the cell cycle. In early G1 phase (pre-start), Whi5 is concentrated in the nucleus (green) and polarity factors are dispersed. As CDK activation occurs, Whi5 is exported from the nucleus (when 50% of Whi5 has been exported the cells commit to enter the cell cycle at “start”). CDK activation triggers localization of polarity factors to a cortical site (red: polarization) from which the bud later emerges (bud). **(B)** Inverted maximum projection montages of selected timepoints for representative cells from movies of wild-type (WT: DLY19654) or mutant (gic1Δ gic2Δ: DLY20961) cells progressing through the cell cycle at 37°C. The cells express Whi5-GFP (top row) and Bem1-tdTomato (bottom row) probes. Cells were synchronized in G1 by pheromone arrest-release, and time relative to start is indicated. Scale bar, 5 μm. **(C)** Display as for **(B)** but with strains expressing Whi5-tdTomato (top row) and GFP-Cdc42 (bottom row). Wild type: DLY21726. gic1Δ gic2Δ: DLY21728. For both sets of strains, wild-type cells polarized shortly after start, whereas gic1Δ gic2Δ cells either failed to polarize (cell 1) or polarized after a delay (cell 2).

**Figure 2:**
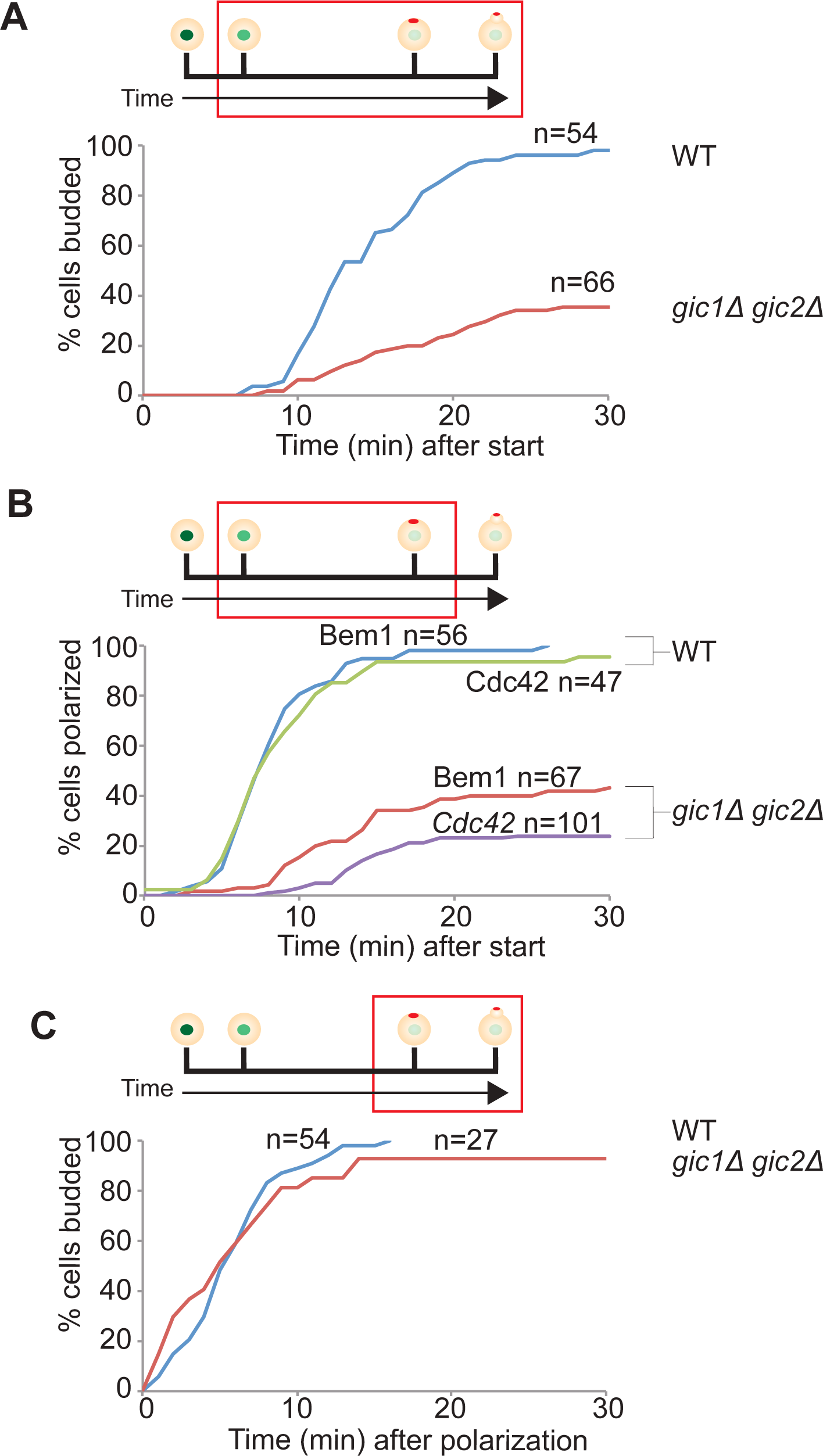
Quantification of polarity establishment in gic1Δ gic2Δ mutants at 37°C. Time intervals between start and bud emergence **(A)**, between start and polarization **(B)**, and between polarization and bud emergence **(C)** were scored from the time-lapse movies described in Fig. 1. Top: schematics as in Fig. 1A, indicating the interval scored (red box). Bottom: graphs showing the cumulative % of cells (y axis) that completed the interval by the indicated time (x axis). The number of cells scored for each plot is indicated (n). **(A-C)** plot data for strains expressing Bem1-tdTomato (as in Fig. 1B), while **(B)** additionally plots data for strains expressing GFP-Cdc42 (as in Fig. 1C). For **(C)**, we only scored the subset of gic1Δ gic2Δ cells that polarized (hence the lower n).

Polarity establishment requires activation of Cdc42, which is promoted by the GEF Cdc24 and antagonized by the GAPs Bem2, Bem3, Rga1, and Rga2 (Howell and Lew, 2012). Thus, one possible basis for the defect in polarity establishment in *gic1*Δ *gic2*Δ cells is that they have insufficient GEF or excess GAP activity. As an initial attempt to test that hypothesis, we compared the abundance of these regulators in wild-type and *gic1*Δ *gic2*Δ cells. We noted no significant differences, either when the cells were grown at 24°C (Fig. 3A) or 37°C (Fig. 3B). Many of the regulators undergo phosphorylation, which is thought to regulate their activity (Bose et al., 2001; Gulli et al., 2000; Knaus et al., 2007; Kuo et al., 2014; Sopko et al., 2007; Wai et al., 2009). Although we detected altered-mobility species in many of the blots, we did not find any systematic difference between wild-type and *gic1*Δ *gic2*Δ cells.

**Figure 3:**
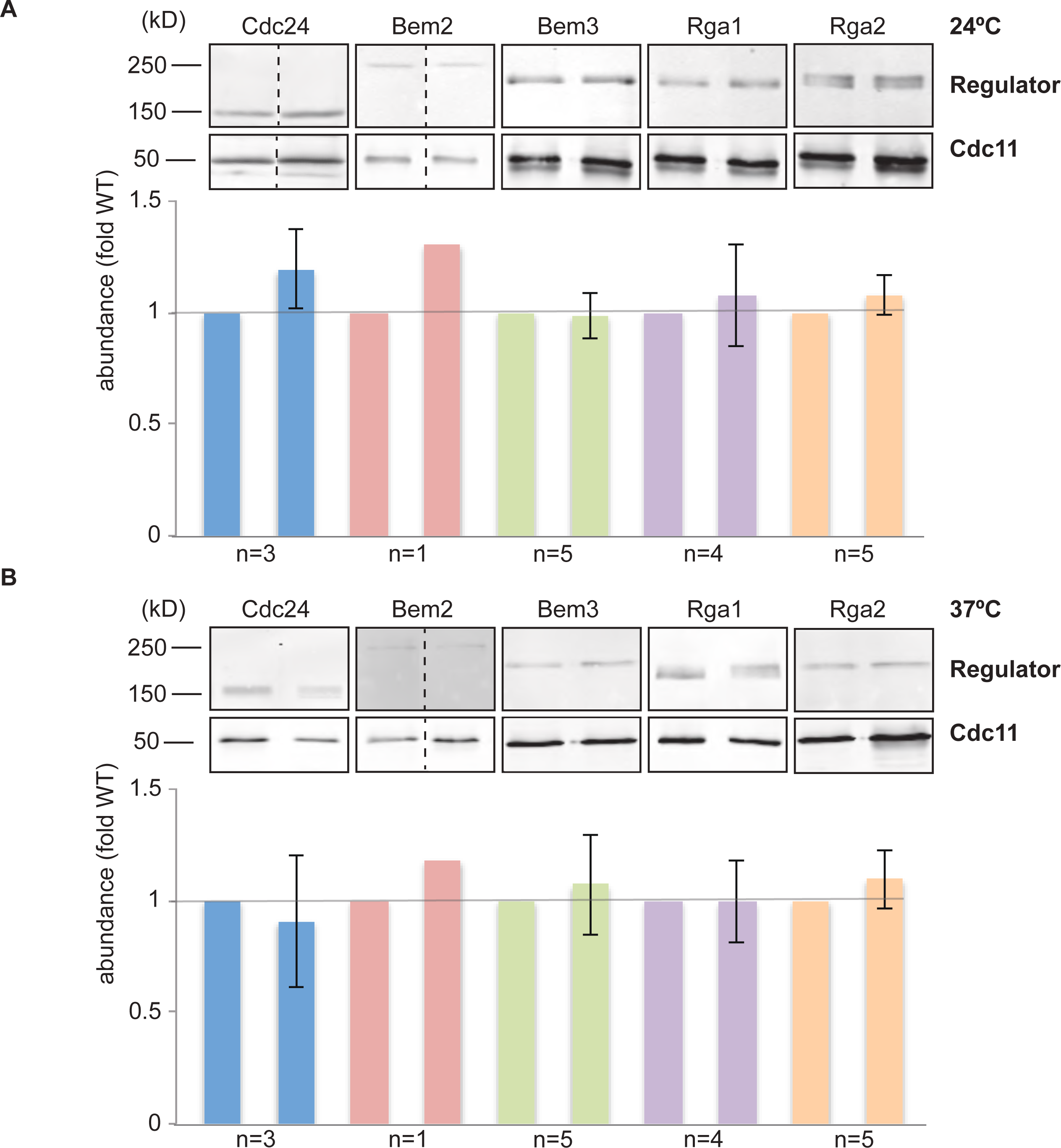
Abundance of Cdc42-directed GEF and GAPs in gic1Δ gic2Δ mutants. **(A)** Anti-HA Western blot to compare the abundance of Cdc24-3HA expressed at the endogenous locus in wild-type (DLY15429) and gic1Δ gic2Δ (DLY21815) strains (left). Anti-myc Western blots to compare abundance of Bem2-12myc (wild-type, DLY8228; gic1Δ gic2Δ, DLY22229), Bem3-12myc (wild-type, DLY11483; gic1Δ gic2Δ, DLY22232), Rga1-12myc (wild-type, DLY21093; gic1Δ gic2Δ, DLY22235), and Rga2-12myc (wild-type, DLY11847; gic1Δ gic2Δ, DLY22232) expressed at the endogenous loci. Loading control is a blot of Cdc11 (a septin) in the same lysates. Cells were grown to mid-log phase and lysates were prepared as described in Methods. Quantification of each blot (fluorescence intensity of secondary antibody for each regulator normalized to its corresponding loading control) is shown in the bar graph below each blot. When independent Western blots were performed, the number of blots is indicated and the bar graphs show mean and standard error of the mean. **(B)** Western blots were repeated using lysates from cells that were shifted to 37°C for 6 h prior to lysate preparation.

We next attempted a genetic approach to test whether mutations in regulators might enhance or suppress the phenotype of *gic1*Δ *gic2*Δ cells. Although *gic1*Δ *gic2*Δ mutants are lethal at 37°C following tetrad dissection (Fig. 4A), this approach was thwarted by a high frequency of spontaneous suppression of the lethality. We speculated that because a subpopulation of mutant cells was able to bud (Fig. 1 and 2), strong selection pressure could be applied to the expanding population, yielding a high spontaneous suppression frequency. Such suppression might occur at many loci or just a few, and we reasoned that in the latter case, identification of the basis for spontaneous suppression might be informative with regard to the specific molecular defect that prevents polarization of a majority of *gic1*Δ *gic2*Δ cells.

**Figure 4:**
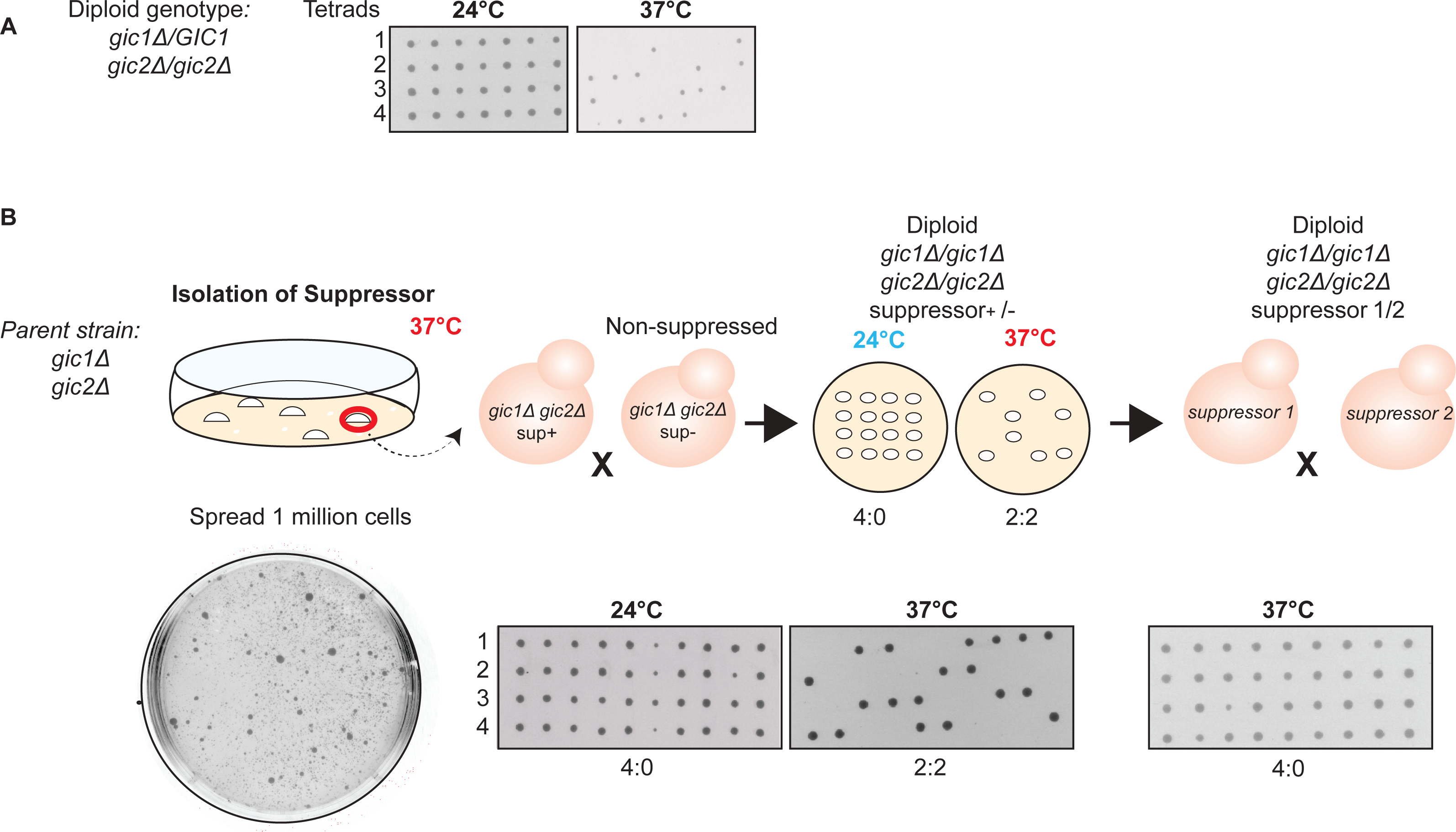
gic1Δ gic2Δ mutants spontaneously acquire a Mendelian suppressor mutation. **(A)** gic1Δ gic2Δ mutants are inviable at 37°C. A diploid strain with the indicated genotype (DLY21711) was sporulated and tetrads (four spores in a vertical column) were dissected onto plates that were incubated at the indicated temperature. Tetrads contain two GIC1 gic2Δ spores and two gic1Δ gic2Δ spores. At 24°C all four spores were viable and gave rise to colonies, but at 37°C two spores from each tetrad died. Replica plating confirmed that the dead spores were the gic1Δ gic2Δ cells. **(B)** Isolation and genetic characterization of spontaneous gic1Δ gic2Δ suppressors. Cells of a gic1Δ gic2Δ strain (DLY19654) were streaked for single colonies. One million cells from each colony were plated on rich media and incubated at 37°C for 3 days. Although most cells died, several heterogeneously sized colonies were able to grow (example plate, bottom left), and one large colony from each independent plate was picked for further analysis. Suppressed cells were mated to a non-suppressed gic1Δ gic2Δ strain of opposite mating type (DLY21941), and the resulting diploids were sporulated and dissected as in **(A)**. Tetrads showed 2:2 viability (middle panels) at 37°C indicating segregation of the suppressor as a single Mendelian locus. Independent suppressed strains (from different initial colonies) were then mated to each other and the resulting diploids were sporulated and dissected as in **(A)**. All tetrads showed 4:0 viability at 37°C (right panels) indicating that the suppressors all map to the same locus. Sequencing confirmed that suppressed strains retained the gic1Δ and gic2Δ mutations.

We picked 10 independent unsuppressed haploid *gic1*Δ *gic2*Δ colonies growing at 24°C, and spread a million cells of each colony on a rich media plate that was incubated at 37°C. Multiple colonies arose spontaneously on each plate, ranging from large to tiny in size (Fig. 4B). We picked a large colony from each plate, and mated them to an unsuppressed *gic1*Δ *gic2*Δ of the opposite mating type. Upon sporulation of the resulting diploids, viability at 37°C segregated 2:2 in tetrads in 9 cases, showing that suppression was due to a single Mendelian locus in these independently derived strains (Fig. 4B and Table 1).

**Table 1:** Tetrad analysis of diploids from crosses between suppressed and non-suppressed *gic1*Δ *gic2*Δ strains

| Suppressor | % of tetrads segregating 2:2 for viability at 37°C | Number of tetrads |
| --- | --- | --- |
| 1 | 100 | 22 |
| 2 | 96 | 25 |
| 3 | 92 | 26 |
| 4 | 78 | 18 |
| 5 | 100 | 25 |
| 6 | 78 | 18 |
| 7 | 100 | 24 |
| 8 | 89 | 19 |
| 9 | 95 | 21 |

To assess whether the independent suppressors occurred at the same or different loci, we performed pairwise crosses between *gic1*Δ *gic2*Δ mutants carrying the different suppressors. In all cases, diploids generated by crossing one suppressed strain to another showed 4:0 segregation for viability at 37°C (Fig. 4B). This indicates that all suppressors are tightly linked, and likely to be in the same locus.

To characterize the suppressed phenotype, we performed live cell imaging of suppressed strains carrying Whi5-GFP and Bem1-tdTomato. We found that suppressed strains were similar to wild-type in terms of the efficiency and timing of polarization relative to start (Fig. 5). Thus, suppression is highly effective in restoring the ability to polarize.

**Figure 5:**
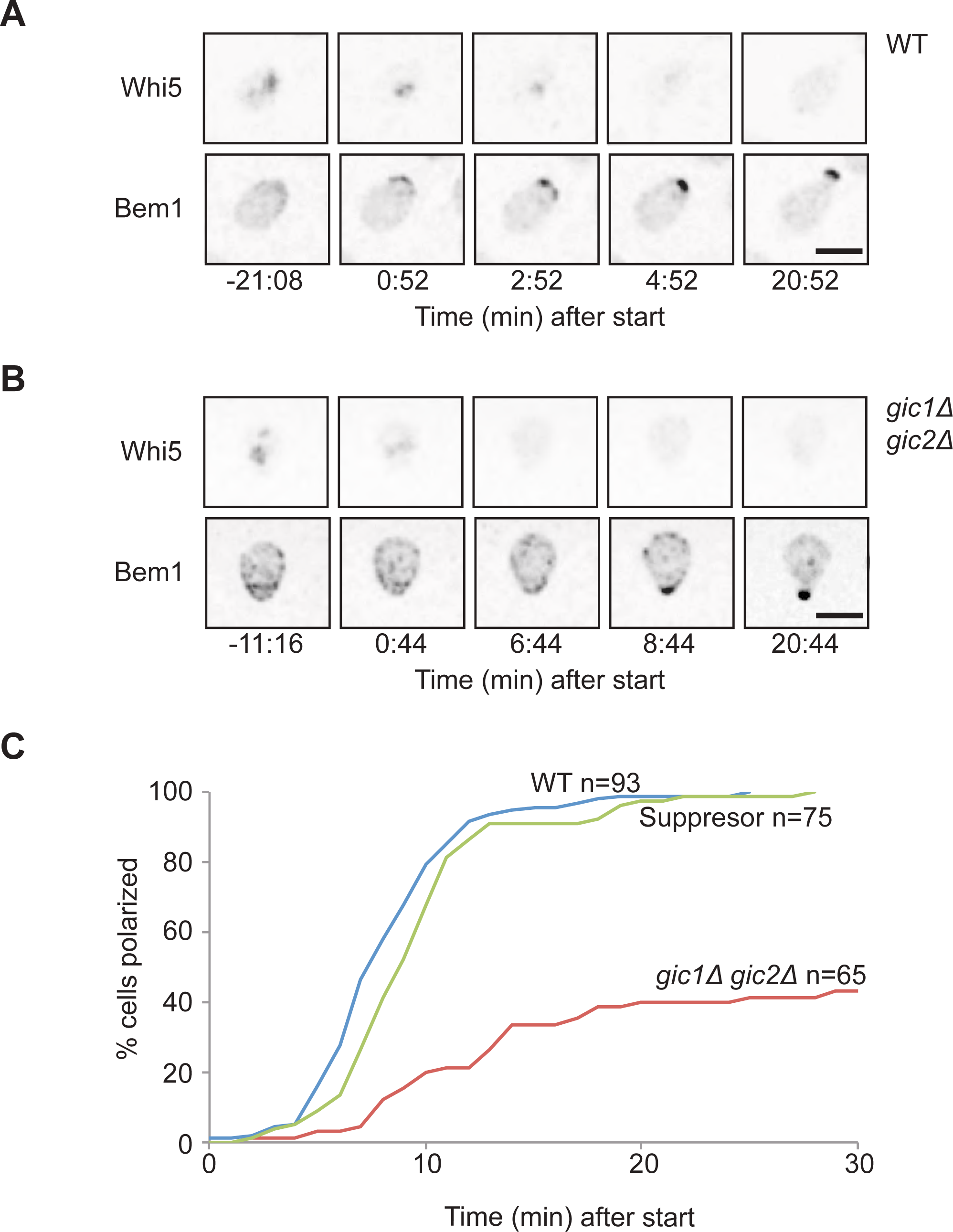
Suppressed gic1Δ gic2Δ mutants polarize like wild-type cells. **(A)** Inverted maximum projection montages of selected timepoints for representative cells from movies of wild-type (WT: DLY19654) or suppressed gic1Δ gic2Δ (DLY22968) cells progressing through the cell cycle at 37°C. The cells express Whi5-GFP (top row) and Bem1-tdTomato (bottom row) probes. Cells were synchronized in G1 by pheromone arrest-release, and time relative to start is indicated. Scale bar, 5 μm. **(B)** Time intervals between start and polarization were scored from the time-lapse movies above as in Fig. 2B. The number of cells scored for each plot is indicated (n). Data for unsuppressed gic1Δ gic2Δ cells is reproduced from Fig. 2 to allow direct comparison to suppressed strain.

## Discussion

Previous studies identified roles for Gic1 and Gic2 in regulating actin or septin organization downstream of Cdc42. As GICs are effectors that bind specifically to GTP-Cdc42, it was natural to expect roles of GICs acting downstream of Cdc42. However, the discovery of synthetic lethality between *rsr1*Δ and *gic1*Δ *gic2*Δ mutants (Kawasaki et al., 2003) indicated that GICs might also act upstream of Cdc42. Our major finding is that GICs are required for efficient and timely polarization of Cdc42 at 37°C, strongly supporting the conclusion that GICs act upstream of Cdc42.

There are several potential explanations for these findings. First, GICs might play dual roles, acting both upstream of Cdc42 and downstream of Cdc42 in separate pathways. A preliminary examination of the levels of known Cdc42 regulators did not reveal any differences between wild-type and *gic1*Δ *gic2*Δ mutant cells, but it remains possible that GICs affect the activity rather than the abundance of these regulators.

Second, GICs may simply act as downstream effectors of Cdc42, mediating cytoskeletal reorganization. Because Cdc42 is known to polarize even in the absence of F-actin (Ayscough et al., 1997; Irazoqui et al., 2003) or polymerized septins (Pringle et al., 1995), this alone would not necessarily yield the observed defects in Cdc42 polarization. However, it could be that the particular cytoskeletal misregulation that occurs in *gic1*Δ *gic2*Δ mutants triggers a stress response that blocks effective Cdc42 polarization. Although stress pathways can act to block polarization (Delley and Hall, 1999; Mutavchiev et al., 2016), we believe this scenario is unlikely.

Third, and perhaps most likely, GICs could operate as part of a positive feedback loop in which GTP-Cdc42 acts to promote further local accumulation of GTP-Cdc42. This would explain why cells lacking GICs have difficulties in polarizing Cdc42, and there is precedent for such feedback in the role of PAKs and Bem1 (Chiou et al., 2017; Kozubowski et al., 2008). However, the mechanism by which GICs might exert such feedback remains mysterious, and given that cells already have one positive feedback pathway it is not immediately obvious why they would require another.

Cells growing at 24°C do not require GICs for successful proliferation, indicating that there are parallel pathways that can operate in the absence of GICs. Moreover, the growth defect of haploid cells lacking GICs can be suppressed by overexpression of Cdc42 (Chen et al., 1997), and diploid cells lacking GICs are able to proliferate successfully even at 37°C (Bi et al., 2000). Other mutants (e.g. lacking the formin Bni1) display more severe phenotypes in diploids than in haploids (Bi et al., 2000). The basis for these differences is unclear. We found that in our strain background, *gic1*Δ *gic2*Δ mutants frequently acquired spontaneous suppressors, and a genetic analysis indicated that several independently isolated suppressors all mapped to the same locus. Identification of the suppressor gene may provide insight into the role of GICs in promoting Cdc42 polarization.

## Materials and Methods

### Yeast strains and growth conditions

The yeast strains used in this study are in the YEF473 strain background (*his3-Δ200 leu2-Δ1 lys2-801 trp1-Δ63 ura3-52*)(Bi and Pringle, 1996) and are listed in Table 2. Standard yeast molecular and genetic manipulations were used to construct strains, with additional precautions due to the high propensity of strains lacking GICs to become genetically suppressed. *GIC1* and *GIC2* deletions were generated by the one-step PCR-based method (Baudin et al., 1993) with pRS304 as template for *gic1∷TRP1* and pRS403 as template for *gic2∷HIS3*.

**Table 2:**
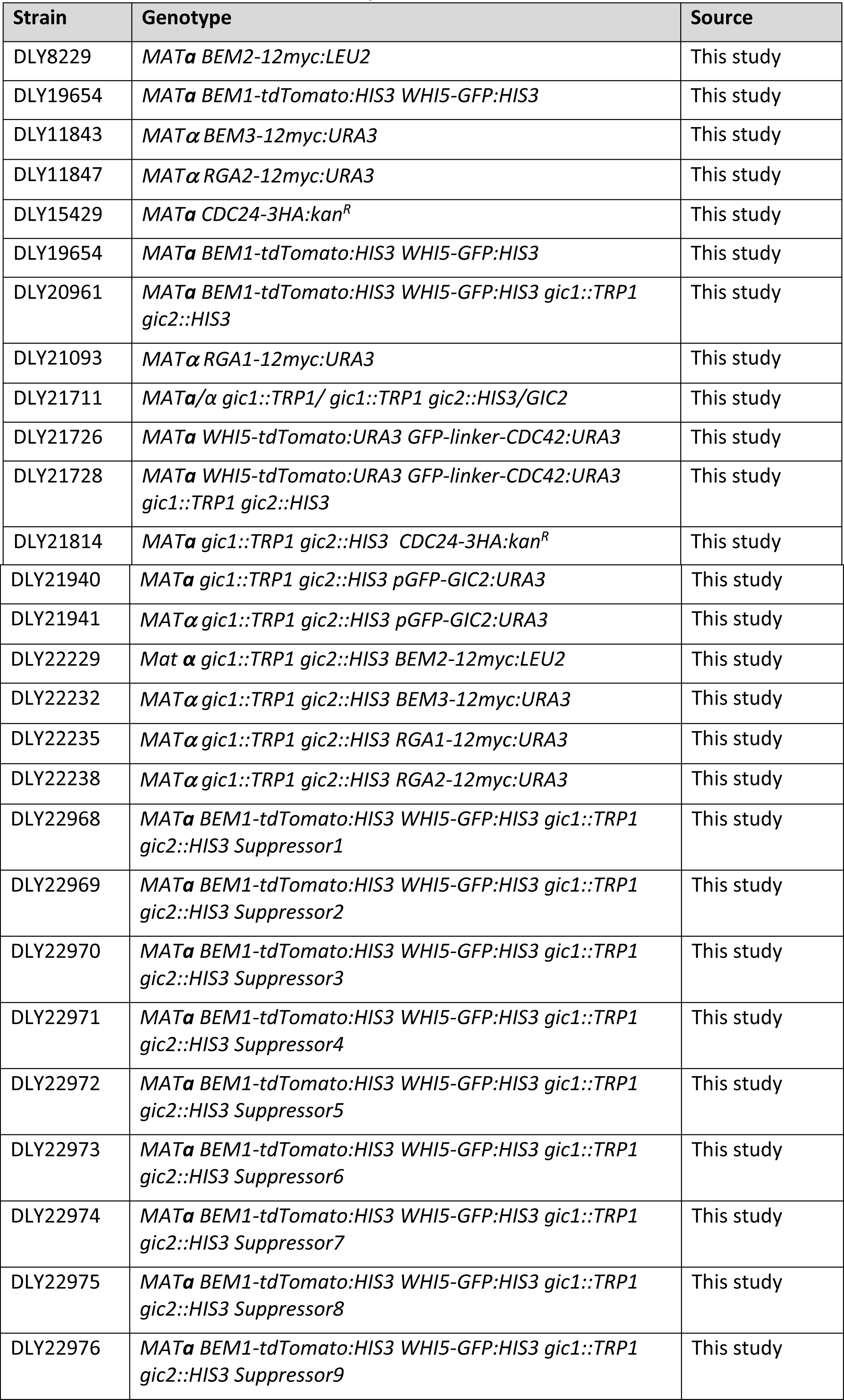
Yeast strains used in this study

Deletions were introduced into diploid strains, and diploids containing at least one wild-type GIC gene were used as strain construction intermediates to avoid selection for suppressors. In cases where strain construction involved a haploid *gic1∷TRP1 gic2∷HIS3* intermediate, we introduced a *URA3*-marked 2 μm plasmid (pDLB2693) carrying wild-type *GIC2* into the parent diploid strain and maintained the plasmid in the derived haploid so as to avoid selecting for suppressors. Loss of the plasmid was induced when needed by growth on plates containing 5-fluoroorotic acid (5-FOA)(Boeke et al., 1987).

Gene tagging at endogenous loci was previously described for Whi5-GFP (Doncic et al., 2011), Whi5-tdTomato (Liu et al., 2015), Bem1-tdTomato (Howell et al., 2012) and Bem2-12myc (Marquitz et al., 2002). The GFP-linker-Cdc42 probe was expressed in addition to endogenous untagged CDC42 (as the GFP-tagged version is not fully functional on its own) as described (Kuo et al., 2014).

The Cdc24-3HA allele was generated by the PCR-based gene modification method (Longtine et al., 1998). Briefly, primers with 50 bp of *CDC24* C-terminus and 3’UTR homology were used to amplify the pFA6 3HA *kanMX* cassette. The PCR product was purified and transformed to tag *CDC24* via standard transformation methods. Proper integration was confirmed by PCR and sequencing.

The Bem3-12myc, Rga1-12myc, and Rga2-12myc constructs were made by cloning PCR products encoding the C-termini of the proteins into a pRS306-based integrating plasmid (pSWE1-myc) containing 12 myc tags and the *SWE1* terminator (McMillan et al., 1998). Digestion at a site within the gene was used to target integration of the plasmid at the endogenous loci, and proper integration was confirmed by PCR checks.

Cells were grown on rich YEPD media (1% yeast extract, 2% peptone, 2% dextrose) or complete synthetic medium (CSM; MP Biomedicals) with 2% dextrose at 24°C as described below.

### Isolation and Analysis of Suppressor Mutants

The MAT**a** *gic1Δ gic*2Δ strain DLY20961 was streaked for single colonies on YEPD plates at 24°C. 10 colonies were picked using sterile toothpicks, and the cells were resuspended in 1 mL of sterile distilled water and counted. 1 million cells from each colony were spread onto individual YEPD plates. Each plate was incubated at 37°C for 3 days, after which plates displayed growth of numerous microcolonies and a few large colonies. A large colony was picked from each of the 10 plates, and mated to non-suppressed MATα *gic1Δ gic*2Δ strain DLY21941. Mating was conducted by mixing cells on a YEPD plate, with a large excess of the MAT**a** strain, so that most MATα cells would mate. Cells from the mating mix were spread on YEPD plates containing 2 μM α-factor to arrest unmated MAT**a** cells, and colonies were tested to determine whether they could sporulate (indicating successful diploid formation) when transferred to 2% Potassium Acetate plates and allowed to grow for 5-7 days.

Asci from sporulating diploids were digested by treatment with lyticase for 5 min. Tetrads were diluted in sterile distilled water, spread on YEPD plates, and dissected with a micromanipulator. Tetrad plates were incubated at 24°C or 37°C as indicated for 3 days. Images of plates were taken on day 3. Spore colonies were replica plated to relevant selective media plates to test for auxotrophic markers. Suppressor strains were then crossed to each other and tetrads were analyzed using a similar procedure.

### Cell synchronization

For imaging experiments, the cells were first synchronized by G1 arrest/release. MAT**a** cells were grown overnight at 24°C in CSM+D, adjusted to 1.5×10^7^ cells/mL, and treated with 2 μM α-factor (Genesee Scientific) at 24°C for 3 h. G1-arrested cells were released from arrest by washing two times with fresh medium, and placed on microscope slabs at 37°C for imaging.

### Microscopy and image analysis

Cells were mounted on a 250 μL slab solidified with 2% agarose on a microscope slide. After putting a cover slip on top, the edges were sealed with petroleum jelly to prevent evaporation. Image acquisition was done using an Andor XD Revolution spinning-disk confocal microscope (Olympus) with a Yokogawa CSU-X1 5000 r.p.m. disk unit, and a 100x/1.4 UPlanSApo oil-immersion objective controlled by MetaMorph software (Universal Imaging). The microscope is enclosed in a temperature-controlled chamber that was set to 37°C 1 h prior to imaging. Fluorophores were excited with 488 nm and 561 nm diode lasers. Images (stacks of 17 z planes spaced 0.5 μm apart) were collected at 1 min intervals using a iXon3 897 EM-CCD camera with 1.2x auxiliary magnification (Andor Technology). Laser power was set to 10% maximum output to reduce phototoxicity. Exposure time was 200 ms for each image. An EM-Gain setting of 200 was used for the EM-CCD camera.

Collected images were deconvolved using Hyugens Essential software (Scientific Volume Imaging). Images were then processed using ImageJ (National Institutes of Health). Z-stacks were collapsed into maximum projection images. Polarization was scored by eye as the first detection of a cluster of the polarity probe (GFP-Cdc42 or Bem1-tdTomato). Whi5 nuclear export was scored using a custom MATLAB graphical user interface (GUI; NucTrackV3.3) as described (Lai et al., 2018). This tool allows for designation and tracking of a region of interest at specific times of interest during the course of the time-lapse. For our purposes, regions of interest were individual cells. The coefficient of variation of Whi5 signal intensity between pixels in each cell was measured and used to determine the time point at which 50% of Whi5 exited the nucleus. Calculated values are normalized to peak intensity for each track. This tool is available upon request from Dennis Tsygankov.

### Immunoblotting

Cells were grown overnight in YEPD at 24°C, and where indicated shifted to 37°C for 6 h prior to harvesting. Cell pellets (about 10^7^ cells) were resuspended in 225 μL cold Pronase buffer (25 mM Tris-HCl, pH 7.5, 1.4 M Sorbitol, 20 mM NaN^3^, 2 mM MgCl^2^) and 48 μL of 100% TCA. Pellet-buffer mixture was stored frozen at −80°C. Once thawed on ice, cells were lysed by vortexing with 280 μL of sterile acid-washed glass beads at 4°C for 10 min. Beads were washed twice with 5% TCA. Lysate was collected and precipitated proteins were pelleted by centrifugation at maximum speed in an Eppendorf centrifuge for 10 min at 4°C. Pellets were solubilized in Thorner buffer (40 mM Tris-HCl, pH 6.8, 8 M Urea, 5% SDS, 143 mM β-mercaptoethanol, 0.1 mM EDTA, 0.4 mg/ml Bromophenol Blue). 2 M Tris base was used to adjust the pH to 8. Samples were heated at 42°C for 3 min prior to loading on a 10% Acrylamide/Bis gel and run for 1 h at 40 mA. Following transfer, membranes were probed with anti-cMyc, or anti-HA (12CA5) (Roche) monoclonal antibodies and anti-Cdc11 polyclonal antibodies (Santa Cruz Biotechnologies) used at 1:1000 and 1:2000 dilution respectively. Secondary antibodies IRDye800-conjugated anti-mouse IgG (Rockland Immunochemicals) and Alexafluor680-conjugated goat anti-rabbit IgG (Invitrogen) were used at 1:10,000 dilution. After washing, Western blots were visualized using the ODYSSEY imaging system (Li-COR Biosciences).

Western blot quantification was done using ImageJ to measure band intensity in individual color channels. Mutant and wild-type bands were always compared from the same blot using lanes with comparable Cdc11 loading controls. After dividing by the loading controls, bands were normalized to the wild-type signal.

## Acknowledgements

This work was funded by NIH/NIGMS grants GM62300 and GM122488 to D.J.L.

